# A new improved and extended version of the multicell bacterial simulator gro

**DOI:** 10.1101/097444

**Authors:** Martín Gutiérrez, Paula Gregorio-Godoy, Guillermo Pérez del Pulgar, Luis Muñoz, Sandra Sáez, Alfonso Rodríguez-Patón

**Affiliations:** Departamento de Inteligencia Artificial, ETSII, Universidad Politécnica de Madrid, Madrid, Spain; Escuela de Informática y Telecomunicaciones, Universidad Diego Portales, Santiago, Chile

**Keywords:** Individual based Model, synthetic biology, cell-cell communication, cell shoving algorithm, ecology, digital proteins

## Abstract

`gro` is a cell programming language developed in Klavins Lab for simulating colony growth and cell-cell communication. It is used as a synthetic biology prototyping tool for simulating multicellular biocircuits. In this work, we present several extensions made to `gro` that improve the performance of the simulator, make it easier to use and provide new functionalities. The new version of `gro` is between one and two orders of magnitude faster than the original version. It is able to grow microbial colonies with up to 10^5^ cells in less than 20 minutes. A new library, CellEngine, accelerates the resolution of spatial physical interactions between growing and dividing cells by implementing a new shoving algorithm. A genetic library, CellPro, based on Probabilistic Timed Automata, simulates gene expression dynamics using simplified and easy to compute digital proteins. We also propose a more convenient language specification layer, ProSpec, based on the idea that proteins drive cell behavior. CellNutrient, another library, implements Monod-based growth and nutrient uptake functionalities. The intercellular signaling management was improved and extended in a library called CellSignals. Finally, bacterial conjugation, another local cell-cell communication process, was added to the simulator. To show the versatility and potential outreach of this version of `gro`, we provide studies and novel examples ranging from synthetic biology to evolutionary microbiology. We believe that the upgrades implemented for `gro` have made it into a powerful and fast prototyping tool capable of simulating a large variety of systems and synthetic biology designs.

---

Synthetic biologists need new computational bioCAD tools to assist them during the design and engineering of new biocircuits. Genetic circuit engineering is moving from single cell devices towards multicellular biocircuits.^1^ Ordinary Differential Equations and Gillespie algorithms are useful tools but mainly for intracellular calculations^2,3^. Computational tools are needed to rapidly predict the dynamical behavior of programmed multicellular microbial communities.

Partial differential equations and individual-based models (IbMs)^4,5^ are the most commonly used models to study cell colonies behavior. Partial differential equations aim to reveal the general dynamics of a population as a whole following global equations.^6^ Alternatively, IbMs derive the global dynamics of the population from local interactions rules described for every single individual (*i.e.*, cell). The behavior of the population then emerges as a result of the interaction between these individuals. IbMs are better suited for simulating multicellular systems such as cell colonies or tissues^4^ as more complex behaviors can be achieved. Because of that, several IbM simulators for cell colonies have been developed. These simulators range from general-purpose and computer-consuming frameworks like CellModeller,^7^ IDynoMiCS,^8^ Biocellion,^9^ CeCe^10^ or `gro,`^11^ to more focalized representations oriented towards studying specific aspects of colonies, such as BSim,^12^ BactoSim,^13^ an IPS by Krone et al.,^14^ COMETS^15^ or DiSCUS.^16^ Other solutions include programming frameworks for constructing IbM simulators like REPAST,^17^ FLAME,^18^ Netlogo^19^ and Mason.^20^

We have chosen `gro`^11,21^ as a base platform to extend because of its flexible architecture and open-source nature. It is a 2D simulator that reproduces microcolony growth within a simple graphical interface, which is ideal for visualizing spatial arrangements. Cell behavior specifications are implemented through its own flexible functional programming language. `gro` was used for simulating synthetic collective intelligence^22^ and studying synthetic bacterial architectures using morphogenetic engineering.^23^ `gro` has also been used at the 2016 iGem by the award-winning Imperial College team to simulate multicellular behavior of RNA circuits in their ecolibrium project.

Despite being a solid simulation platform, we identified some shortcomings of `gro` as a prototyping tool:

- The algorithm for handling cell mechanics in the colony is too costly and limits the number of cells that can be simulated: at about 3000 cells, the simulator’s execution speed greatly slows down.
- Specifying genetic circuits in `gro` requires the abstraction of its components to align with `gro`’s functional programming language.

Accelerating and enriching `gro` would make it an an excellent tool for executing multicell modeling and simulations. This was our motivation for improving `gro` and adding the new functionalities described in this paper. The flexible architecture of `gro` allows for features to be coupled easily. We have tackled the presented limitations through the following improvements: We constructed and integrated a new physics engine, CellEngine. It was developed to improve the performance of simulations involving a large number of cells (in the order of 10^5^ cells) through the use of a new algorithm. We developed a Probabilistic Timed Automata based library, CellPro, that encapsulates and simulates gene expression with simplified and easy to compute digital (bearing values of 0 or 1) proteins with a noisy behavior based on probabilities. We also built a new specification language layer, ProSpec, to define genetic circuits in terms of digital proteins. ProSpec also handles cell behavior by evaluating logical conditions in terms of digital protein values.

In addition, we have implemented a set of other features to further improve the simulator. First, we implemented a new library called CellNutrient that handles nutrient uptake and its influence on growth. We also built a new library that extends intercellular and environmental signaling capabilities called CellSignals. Finally, we added another form of intercellular communication: bacterial conjugation.

Our new version of `gro` can simulate larger bacterial colonies in a shorter time than the original version of the simulator (see Figure 1). Starting from a single bacterium, colonies of 10^5^ simulated bacteria are now reached in around 20 minutes in a standard computer (see Figure S2). The original version of `gro` reaches about 5000 bacteria in the same time. For the original version to simulate 10^5^ bacteria, it would require over five days of computation.^24^ The improved `gro` also proved to be faster than other powerful simulators such as CellModeller^7^ by about an order of magnitude (see Figure S2).

**Figure 1:**
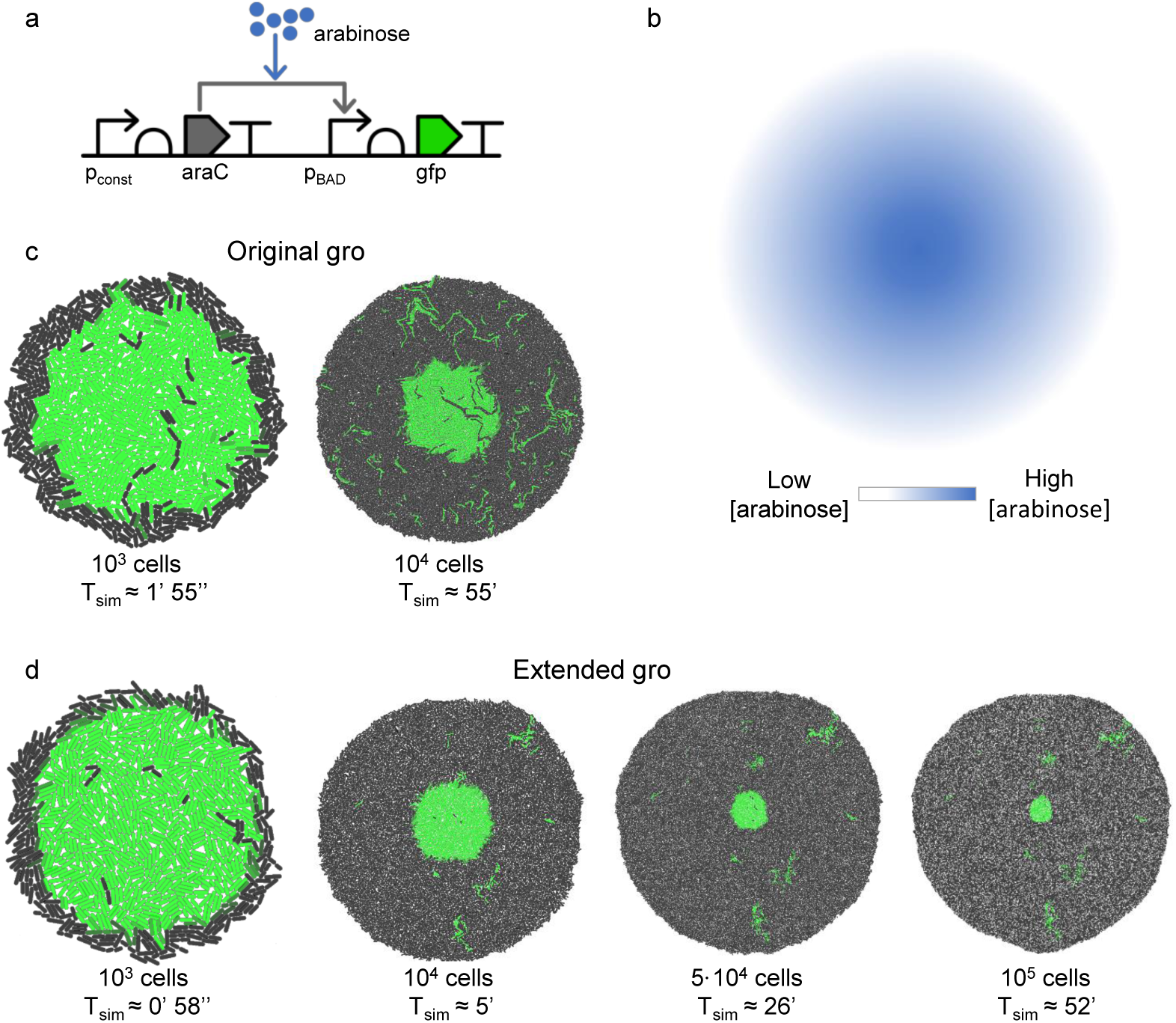
Visual performance comparison of the original `gro` and our extended version. The new version of `gro` simulates larger bacterial colonies. We compared the execution speed and size of the colonies for the simulation of an arabinose reporting circuit. a) Genetic circuit. When arabinose is present in the environment and sensed by the circuit, it is reported by GFP. b) A radial gradient of arabinose serves as the spatial cue for expressing GFP. Therefore, the expression of GFP is expected to be located at the center of the colony. c) The original version of `gro` is able to simulate a colony of about 10^4^ cells carrying and executing this circuit in an hour. d) Our version reaches over ten times more cells in the same time. The 10^4^ bacteria simulated by the original `gro` are reached by our version in a little over 5 minutes.

Each of the extensions will be described in detail throughout the course of this paper. We complement the description of our extensions by presenting some relevant case studies and synthetic circuit examples to illustrate the new functionalities of the new version of `gro`.

## Results and discussion

`gro` was originally conceived as a simulator that relies on three major components:

1. Chipmunk^25^, an open-source physics engine that handles the mechanics of how bacteria are placed and move in the environment when they grow and divide.
2. CCl,^26^ a guarded command based language for programming and modeling control systems. This language is used to specify cell behavior and control the environment in which colonies grow.
3. `gro`, the biological backbone of the simulator. Cell dynamics and behavior are implemented in this subsystem.

The goal of this work is to extend the capabilities of `gro` for it to be a more realistic and complete tool for the simulation of synthetic microbial systems. The results are presented as either brand-new components, extensions or replacements for the components of the original version of the simulator. These improvements allow for new simulations that were not previously feasible, or required complicated programming to be executed.

Most of the presented extensions (CellEngine, CellPro, CellNutrient and CellSignals) are provided as standalone external libraries that couple with `gro`. CellEngine acts as a replacement for Chipmunk, while the rest of the libraries add new features. It should be noted that all these standalone libraries can be used by any other software. The protein oriented specification language ProSpec was built as an extension to the CCL language. Finally, the conjugation method was implemented as a property of plasmids triggered by proteins. All these extensions integrate into the main workflow of `gro` as shown in Figure 2.

**Figure 2:**
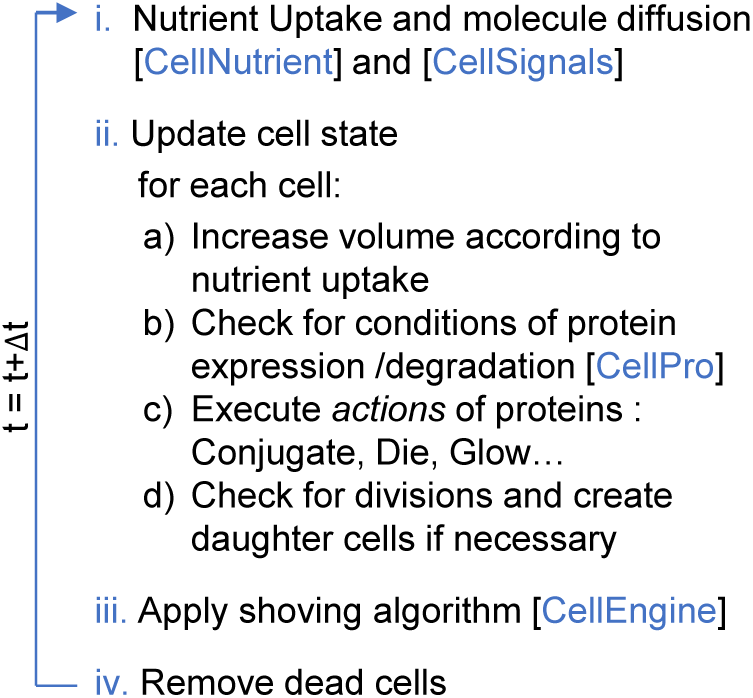
New `gro` execution workflow. This workflow summarizes the update steps that the simulator applies to each cell in the colony and the environment at each time step. Each new task is mostly executed by a single library: Cell mechanical interaction - CellEngine, Nutrient uptake - CellNutrient, Gene expression and cell behavior - CellPro, Environmental and intercellular signals - CellSignals.

## CellEngine

We implemented a physics engine, called CellEngine, in C. It is focused on the fast simulation of growing bacterial colonies. Physics computation has been optimized for rigid rod-shaped bodies, like *e. coli* bacteria, in 2D non-inertial systems. Using our new algorithm explained below, a speedup of two orders of magnitude was achieved when compared to the original version of the software.

In a bacterial colony, each cell grows a certain percentage of its size during each time interval. In a physics engine, this small size increment generates an overlap among the bodies, which are the physical representation of cells. The basic workflow to resolve the overlap consists of two stages:^27^ collision detection and collision response. The collision detection stage^28^ identifies overlaps among bacteria and stores contacts with information about the collision, such as the contact point, magnitude and direction. Then, collision response^29^ performs the physical arrangement of the bodies (*i.e.*, cells). In our case, rigid-body dynamics are used for the computation of linear and angular displacement of the bodies (Figure S3).

Typically, after contact resolution and due to bacteria packaging and lack of free space, the collision response stage generates other contacts, which must in turn, get resolved. Therefore, contacts are resolved through an iterative method that finally displaces the overlap outwards of the colony. This type of problem belongs to the many-body problem,^30^ whose direct methods are computed in O(*N*^2^), being *N* the number of bodies involved. The solution proposed by the original `gro` belongs to this computational complexity order. In our case of study, the population grows exponentially, causing simulations to quickly become too computationally costly. The proposed solution presents a novel method able to compute an accurate approximation in O(*N*) for the same problem.

The reduction in computational complexity is based on two fundamental assumptions: First, the location and angular orientation of a bacterium depend mostly on the neighboring bacteria. Second, colonies tend to grow outwards radially.

Within the scope of these two assumptions, we approximate the solution to this problem by identifying two types of forces exerted for any given bacterium (see Figure 3a):

1. *Local forces*, exerted by nearby bacteria within a distance *k*, that dictate the degree of precision for the movement and rotation of the current bacterium.
2. A *global force* that pushes the bacterium outwards of the colony. This force depends on the number of bacteria between the current bacterium and the location of the center of the colony.

**Figure 3:**
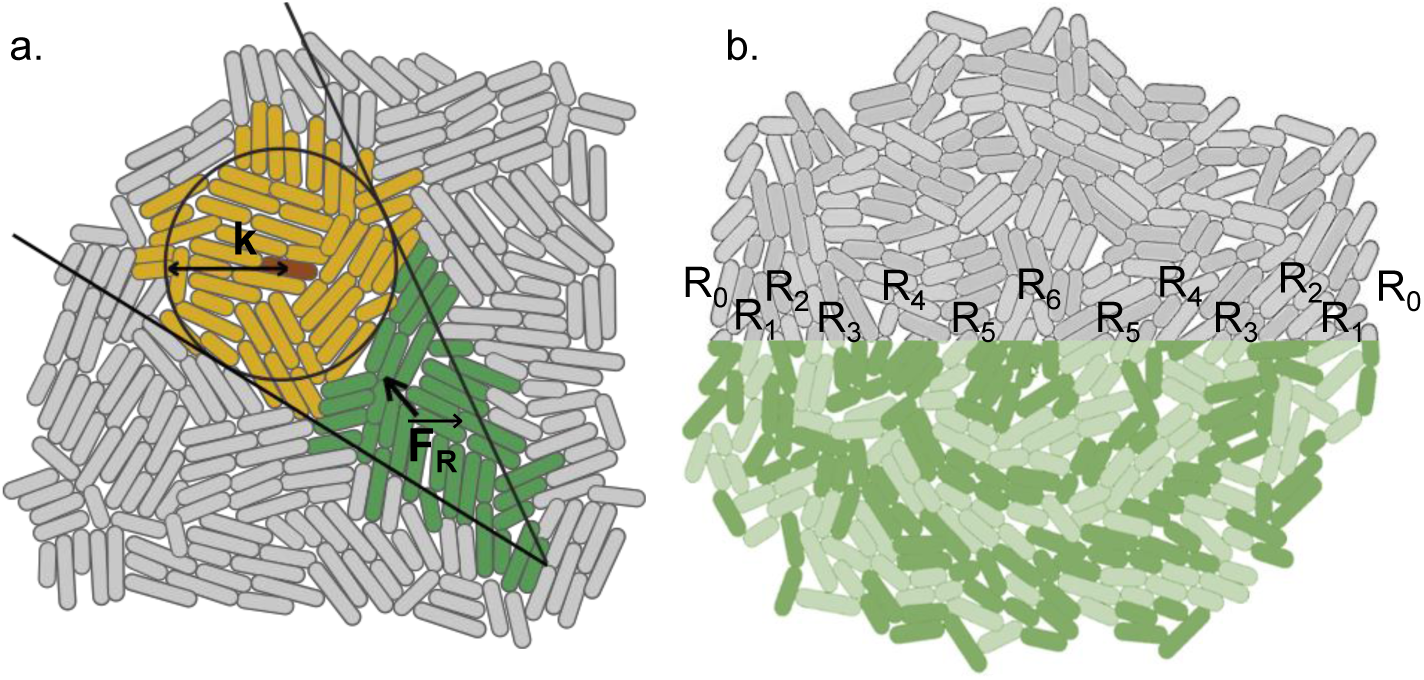
a) Representation of the proposed approximation of colony expansion. The computed forces over each individual bacterium are divided into two types. *Local forces* are generated by the nearby bacteria that exert pressure when growing. *Local forces* are computed only from pressure coming from bacteria that lie within a distance *k* (yellow region). There is also a *global force*, 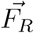 that is exerted radially from the center of the colony. It is a force proportional to the number of cells inwards of the red bacterium (green region). This thrust drives the red bacterium and relocates it at a distance proportional to the module of 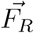. If *k* is kept constant throughout the simulation, the calculation of force and position for each cell is made independent of the colony size, achieving an *O*(*N*) solution. b) Visualization of the ring assignation. Each ring has a width size of a single bacterium. Rings are identifiers of equal pressure bacteria that adapt dynamically as the colony grows. Outer rings experience less pressure than inner rings. The highest pressure of the colony is located in the center ring. Rings act as guides to apply the expansion algorithm.

The proposed approximation is based on the splitting of the colony into concentric rings of one bacterium width. The rings are grouped into sets of width *w* rings that represent the radius *k* of the *local force* action, while the *global force* will be exerted by the bacteria from the inner rings (see Figure 3a). The algorithm transforms the colony into concentric annular sub-colonies of width *w* bodies. When the colony increases in population, the number of rings increases, but the width of every subset, *w*, will remain constant leading to a O(*N*) global solution. Algorithm 1 describes the transfer of overlap in the colony outwards using the rings as guides. It is executed at each time step and has two main stages: Ring tagging and expansion.

The ring tagging stage of the algorithm assigns each bacterium of the colony to a ring (see Figure 3b). It starts from the edge towards the center of the colony. Ring tagging is composed of two phases: edge detection (Algorithm S1) and ring assignment (Algorithm S2).

The second stage of the algorithm is the expansion stage (see Figure 4). It involves the grouping of rings and seeks to relax pressure in the colony. It takes place in an outward manner (from the center of the colony towards the edge) and is composed of two phases: relaxation (Algorithm S3) and relocation (Algorithm S4). The expansion stage selects a set of *w* rings to be grouped and upon which the relaxation phase will be applied. The relaxation phase solves overlap within the set of *w* grouped rings. For each set, bacteria inwards of the set are assumed to have infinite mass, and bacteria outwards of the set are assumed to be non-existent. Taking these constraints into account, a cycle of collision detection and collision response is executed to relieve the pressure of the set. This phase generates overlaps with bacteria in the outward group. To resolve this new overlap, a relocation phase is entered. In this phase, the outer overlapping ring is moved outwards preserving the contact points and relative positions with its neighbors.

**Figure 4:**
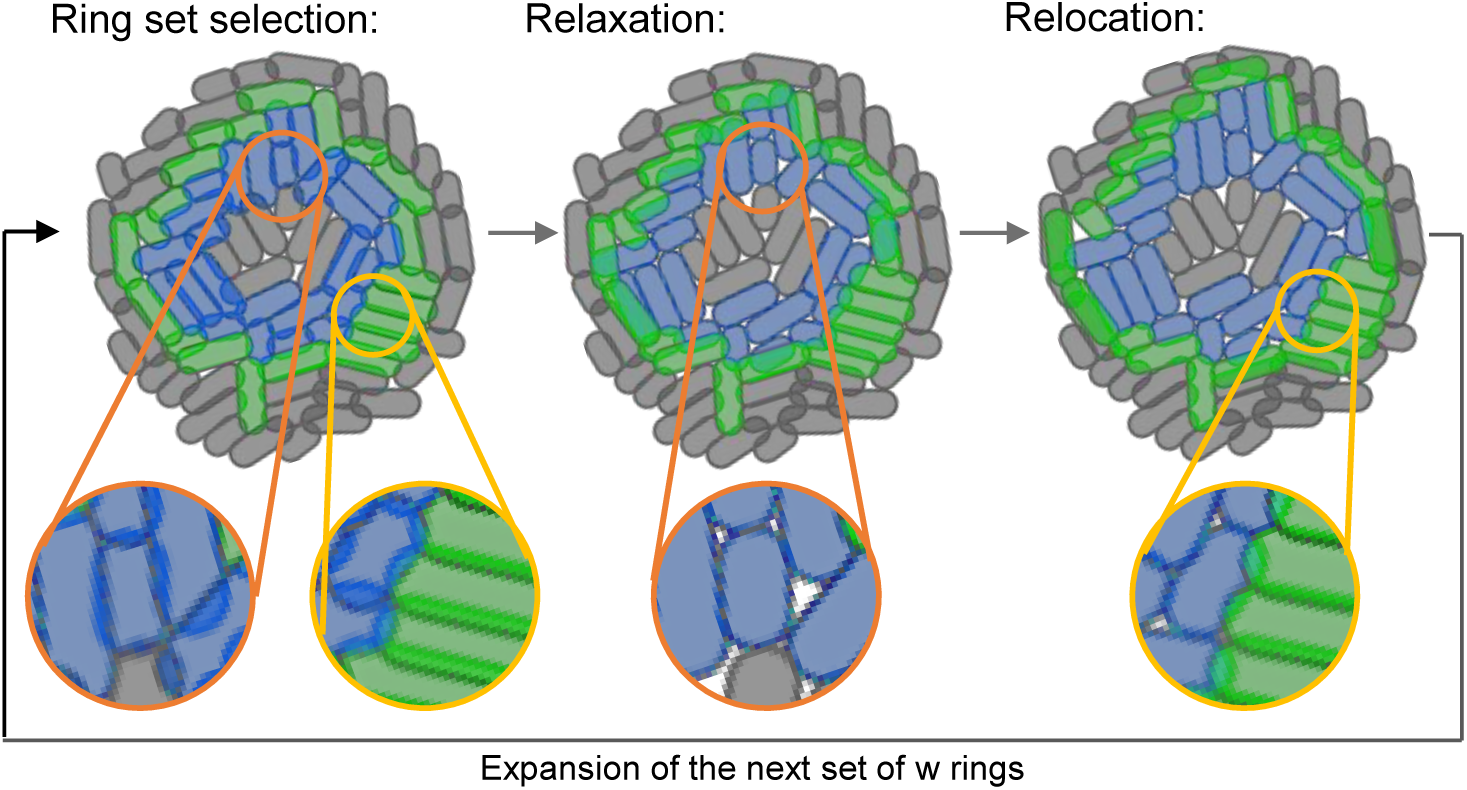
Depiction of the expansion stage. Expansion is performed by, first, relaxing the bodies belonging to the set of blue rings and, later, by relocating bodies belonging to the green ring. The colony on the left shows the organization of bodies before the expansion is performed. At this phase, a set of rings is selected for relaxation and relocation phases. In the central image, the relaxation phase of blue bodies is performed. In this phase, overlaps among blue cells are resolved. Gray cells, inwards of the blue ones, are assumed to have infinite mass. Orange zooms show the overlap resolution at blue-blue (same ring set) and gray-blue (with the inner ring) interfaces performed by the relaxation phase. As a consequence of relaxation, blue bodies dramatically overlap with the outer ring bodies in green. During the relocation phase (right) the green ring bodies are directly re-positioned to maintain the relative position they had with respect to their neighbors before the expansion phase. Yellow zooms shows how the green-blue interface remains invariant thanks to the relocation phase. The whole process is repeated over all subsets of *w* rings in an outwards manner.

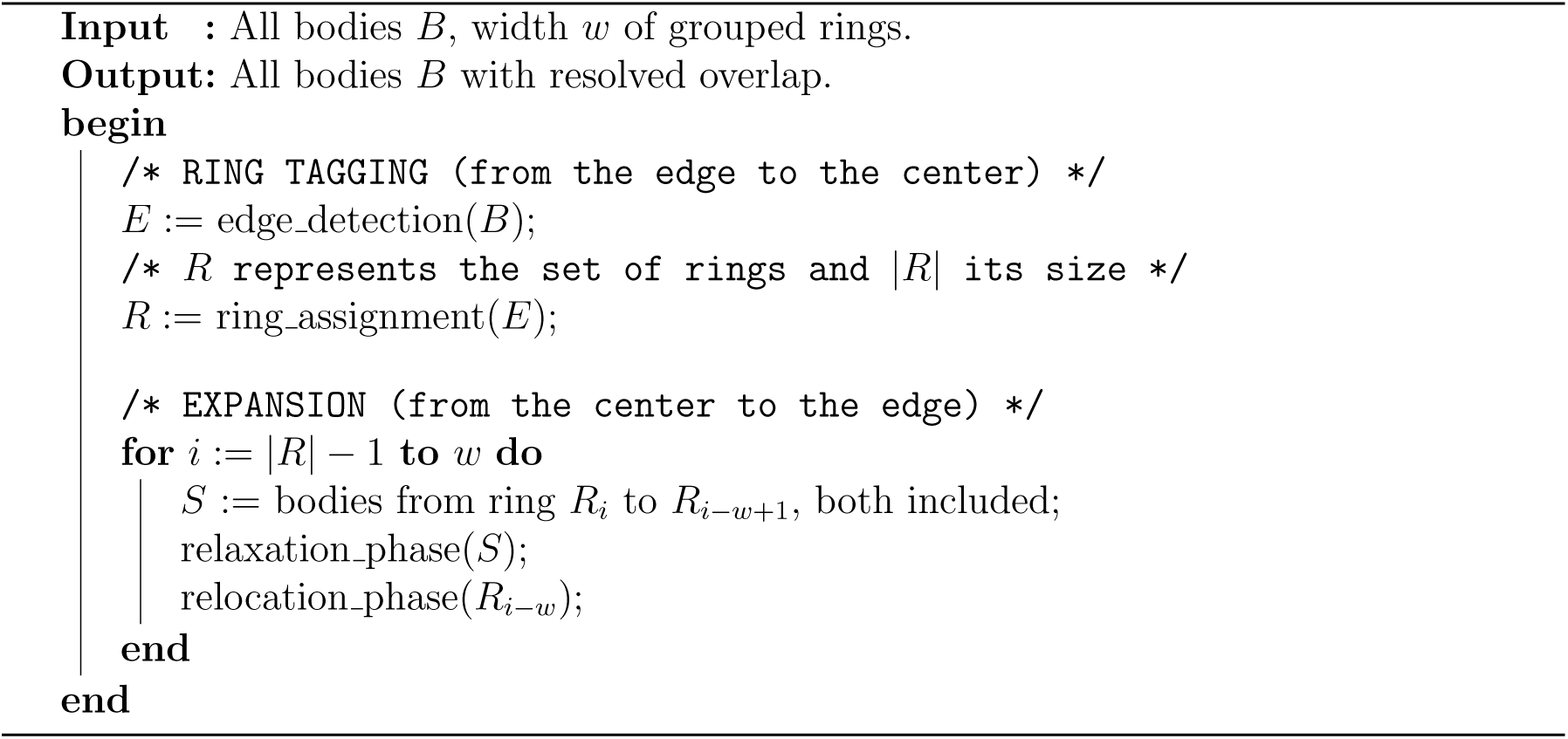

### Algorithm 1

Top-level CellEngine algorithm. It involves two stages: ring tagging and expansion. In the ring tagging stage, each body is assigned to a ring, starting from the edge towards the center of the colony. In the expansion stage, the selection of each group *S* (formed by *w* contiguous rings) will shift outwards one ring at a time after each iteration of the relaxation/relocation phases (see Figure 4). This happens until the selected ring group *S* reaches the colony edge. The relaxation phase solves the contacts generated by bacterial growth by applying rigid-body dynamics. The relocation phase displaces the next outer ring from the current set. The native algorithm implemented in `gro` (using Chipmunk) applies the relaxation process to the whole colony, whereas in CellEngine, the relaxation phase is applied separately into grouped ring sets of width *w*.

We contrasted the performance and quality of CellEngine with the original algorithm of `gro` (using Chipmunk). Cross validation showed that no significant differences are observed in terms of quality when using the proposed algorithm (see Figure S1). Still, CellEngine is able to simulate a hundred times more bacteria for the same computation time.

## CellPro

The original version of `gro` defines a functional, rule-based programming language for specifying bacterial and environment behaviors. Rules are codified as guarded commands *G*: *C* in which the block of instructions *C* is only executed if the guard *G* is satisfied. The abstraction of cell behavior into a rule-based paradigm involves, first, the translation of all elements into variables. Then, it is necessary to describe the relation among these variables in terms of rules. These rules will ultimately update the values of the variables, implementing the specific behavior of the cell. This programming paradigm allows for a broad range of simulations to be implemented, since any level of abstraction may be reached. For instance, a cell response to a given signal can be directly programmed on a high level abstraction, or it may be reached through the specific simulation of a genetic network whose product is signal-sensitive.

This freedom of choice and apparent simplicity of the rule-based programming implies a rigorous and case-sensitive analysis for every change in the design of the simulation. This is one of the largest drawbacks for the average use of `gro`, as it scales poorly. Our proposed motivation is to shift towards a more constrained representation in which behavior of the cells emerges from the interactions occurring at the genetic level. Although this alternative is not as open as the original approach, it fits better with how cells interpret environmental and genetic information. Under this paradigm, genetic circuits are defined in terms of interactions among proteins. Specifically, our hypothesis is that bacterial behavior is driven by protein expression levels. Therefore, cell reaction to its environment is given by cell protein levels.

We have developed a library, CellPro, that simulates gene expression dynamics using digital proteins. Proteins are digital entities with two possible states. When the protein is produced, its value is set to true, and when the protein is degraded or is otherwise absent, its value is set to false. Protein state values are unequivocally defined by the evaluation of the promoter that regulates the protein-associated gene. Promoters are represented as Boolean functions (YES, NOT, AND, OR) and their evaluation is given by the state of the transcription factors regulating them.

Suppose a promoter *Prom* regulates a gene *G_i_* whose expression output is protein *P_i_*. If promoter *Prom* is ON for more than *t_act_* minutes, then the expression value of protein *P_i_* goes from 0 to 1. Alternatively, if promoter *Prom* is OFF for more than *t_deg_* minutes, then the value associated to the expression of protein *P_i_* goes from 1 to 0. *t_act_* and *t_deg_* are two characteristic mean times associated to each protein. They follow a normal distribution that captures the cell to cell variability.

CellPro is formed by an interconnected set of binary asymmetric noisy channels^31^ where each channel represents a protein. The asymmetric noise behavior of every channel (*i.e.*, protein) is given by the two probabilities *p*(0|1) and *p*(1|0) of misevaluation associated to each promoter. This can be seen as a mutation or an anomaly in the promoter that makes it insensitive to its transcription factor.

The reader can relate the model here presented to a Probabilistic Timed Automata (PTA)^32,33^. For each time step during the simulation, every bacterium has an associated state given by the set of all of its protein expression values. Figure 5 shows the representation of a GFP reporter protein induced by araC in terms of our proposed PTA.

**Figure 5:**
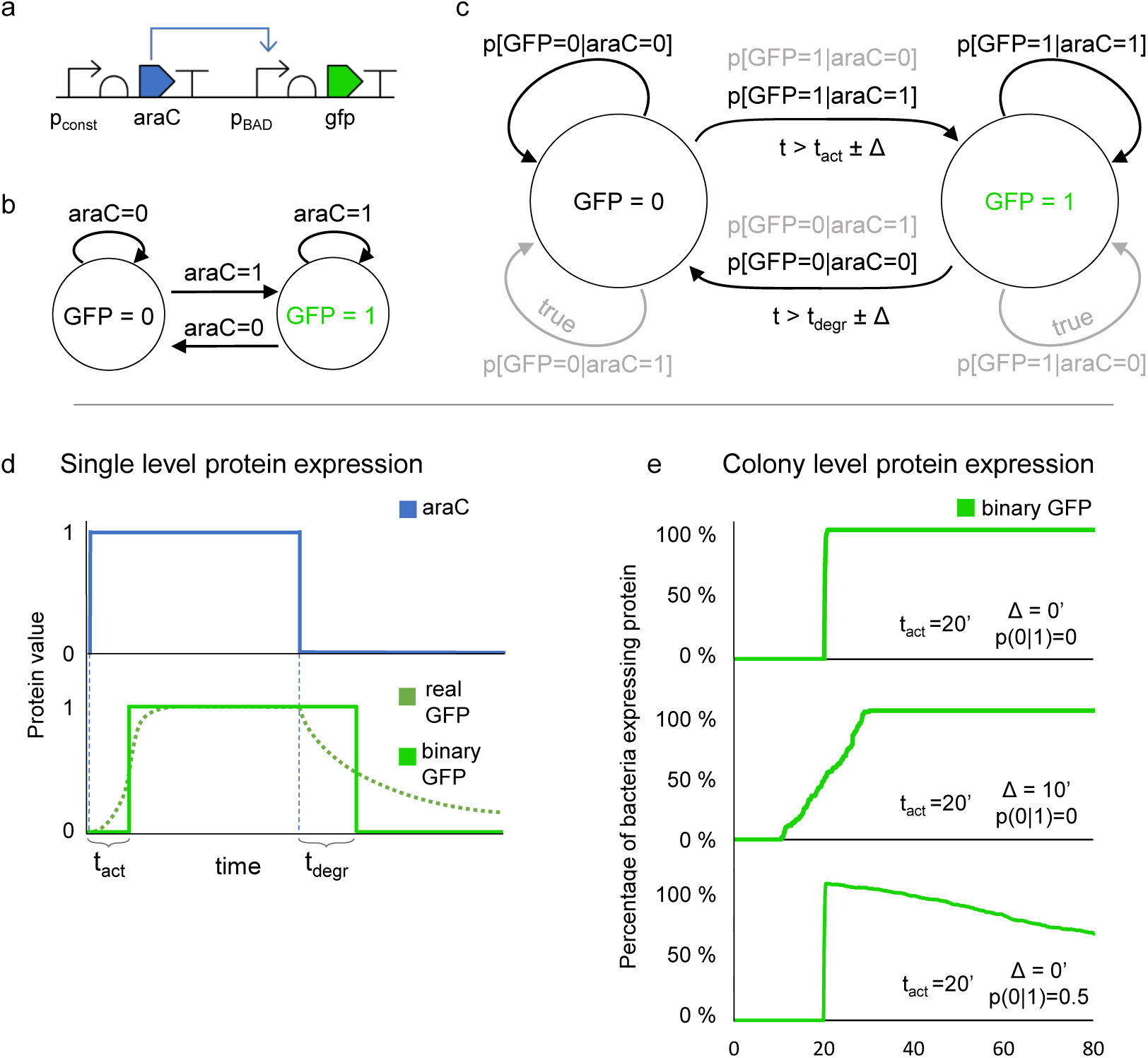
Probabilistic Timed Automata model and simulation data for a YES(araC) gate. a) The output of the circuit (*i.e.*, GFP) is triggered by the value of *araC*. For the sake of simplicity, we assume arabinose to be always present in the environment. b) The dynamics of the circuit, when no noise or delays are present, is represented in the form of an automaton where the state is given by the value of *araC* in the system. c) A detailed version of the dynamics is modeled by a PTA. The delays for transitions between states are dictated by *t_act_*, *t_degr_* and their associated variabilities. Also, noisy behavior can lock the system in the state to where it transitions. Black edges correspond to the transitions shown in subfigure b, while light gray edges are transitions stemming from noise in the system. Both kinds of transitions occur with a certain probability. d) Depiction of the circuit running in a single cell with no noise. *t_act_* and *t*_*deg*_ direct the behavior of the system. e) The circuit running in a colony is shown in different scenarios: i) No variability in the activation time of GFP (20 minutes) and no noise. ii) Variability of 10 minutes in the activation time of GFP and no noise. iii) No variability in the activation time of GFP and an activation noise probability (*i.e.*, p[GFP=0|araC=1]) of 0.5 after 100’.

The entities represented in CellPro are organized hierarchically. Proteins are the fundamental logic unit. Above them, Operons cluster a set of Proteins under the control of a single promoter. Finally, Operons might be distributed in Plasmids, which are at the top of the hierarchy. This hierarchy must be taken into account when a genetic circuit is being specified (see Figure S4).

## ProSpec

As a support for CellPro we have built a new specification language layer: ProSpec. The philosophy behind CellPro and the protein-oriented specification ProSpec is to implement simulations in a way that is easily translatable from a genetic circuit description. Prototyping a circuit as directly as possible helps quickly tinker with and test designs that come from the lab and vice versa (*i.e.*, implement the design with fewer abstraction barriers in the lab). Defining an experiment using ProSpec can be broken down into four parts:

1. Library inclusion, parameter setting and definition of global variables and values.
2. Genetic entities from CellPro and their interactions. *Proteins, Operons* and *Plasmids* are defined in this part.
3. *Actions* associated to the protein state. *Actions* are hard-coded functions that execute a specific function of the cell. The presence or absence of a protein (or of several of them) triggers specific *actions* that direct cell behavior. Examples of these actions are shown in Table 1.
4. Top level programs (main function and other functions) and global control.

Parts one and four have not changed from the original `gro` specification language. However, in part one, we have extended the set of parameters that correspond to the new libraries: CellNutrient and CellSignals. These parameters and their definition are shown in Table S1.

Simulation data can be collected at two different levels: single cell data or aggregated colony data. Functions for collecting data in .csv file format were implemented for both scenarios. In the single cell case, the physical features of the cell and its protein state are recorded at each time step of the simulation. For colony-level data collection, logical conditions depending on protein expression values are stated and evaluated on the colony. Cell counts satisfying these conditions are then dumped into the output file at each time step.

Figure S4 shows a comparison between guarded commands and a ProSpec specification for the simple system presented in Figure 5. Both specifications are equivalent. The guarded command based specification provides a flexible framework for describing an experiment. However, it needs to explicitly handle many aspects of the simulation such as clock update, protein value control and delay calculations. This solution scales poorly for more complex circuits as all of the elements for the simulation must be manually controlled. CellPro and ProSpec encapsulate many of these tasks, and require that the programmer only input core parameters to define the behavior of the genetic circuit. The dynamics of the circuit is then automatically run.

Clauses of our specification language layer may be used in conjunction with the guarded commands of the original `gro` language and do not invalidate them, but on the contrary, are complementary.

## Other libraries: CellNutrient, CellSignals and Bacterial Conjugation

### CellNutrient

`gro` imposes an exponential growth for each bacterium described by Eq. 1:

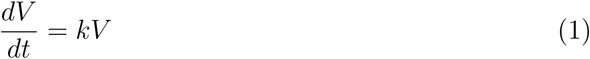

where *V* is the volume of the bacterium and *k* is the growth rate of the bacterium, which was a constant parameter in the original version of `gro`. However, in a petri dish, the growth rate of a given bacterium can be modulated by external factors such as nutrient availability, waste accumulation or even pressure.^34,35^

**Table 1:** Description of *actions* performed by cells in our version of `gro`

| Action | Description | Input parameter |
| --- | --- | --- |
| *Glow* | Change bacterium color | RGB color code |
| *Die* | Removes cell from the simulation |  |
| *Growth rate* | Change growth rate | $k_{max}$ [$\mu m/min$] |
| *Conjugate* | Transfers plasmid to random neighbor bacterium | $\gamma$ [events/generation time] |
| *Lose plasmid* | Remove plasmid from cell | plasmid id |
| *Entry exclusion* | Prevents plasmid conjugation | plasmid id |
| *Read signal* | Check signal at cell location | signal id |
| *Emit signal* | Deposit $E_{max}$ signal at cell location | signal id,<br>$E_{max}$ [$molecules/\Delta t$] |
| *Absorb signal* | Absorbs $A_{max}$ signal at cell location | signal id,<br>$A_{max}$ [$molecules/\Delta t$] |
| *Absorb signal*<br>(*Cross-feeding*) | Absorbs $A_{max}$ signal and modulates growth | signal id,<br>$A_{max}$ [$molecules/dt$],<br>interaction type: (+, -) |
| *Absorb signal*<br>(*Quorum sensing*) | Absorbs $A_{max}$ signal and acts as a protein for CellPro | signal id,<br>$A_{max}$ [$molecules/dt$],<br>signal threshold,<br>protein id |

To simulate a more realistic limitation of growth, we implemented CellNutrient,^36^ a library that offers a built-in Monod-based growth dynamics. If used, CellNutrient modulates the growth rate *k* as a function of nutrient availability in the vicinity following a Monod equation^37^ shown in Eq. 2:

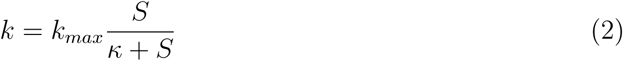

where *k_max_* represents the maximum growth rate, *S* is the available nutrients, and *k* is the half saturation constant.

The consumption rate of the nutrients, their initial amount and their distribution in the environment can be set in the library inclusion part of the experiment specification (see Table S1). Nutrients are consumed by the cells from a grid that stores the amount of nutrient at their specific given location (see Algorithm S6). The grid is dynamic in size, extending its size whenever a bacterium reaches any border of the grid.

A key parameter in the growth dynamics of a single colony is the radial growth. This is the speed of the front of the colony.^38^ If growth is unlimited (as it is implemented in the original version of `gro`), bacteria within the colony grow and divide continually, leading to global exponential growth. This unlimited population growth translates to an exponential radial growth rate. The area of the colony π*R*^2^ increases exponentially, therefore the colony radius *R*, varying proportionally to the square root of the area, increases exponentially as well. However, actual agar plate bacteria do not follow this dynamics. When bacteria from the inner regions of the colony stop growing after some time, the radial growth slows down and becomes linear for long time periods.(*39–41*) Since only the external bacteria divide, the expansion is driven solely by the outer ring of non-exhausted bacteria. Using CellNutrient, we can observe this linear radial growth when the number of bacteria that do not grow is much larger than the number of bacteria still growing (see Figure S5). Other experimentally observed colony morphologies^42^ can also be simulated by tuning the correspondent initial profile of nutrients in the grid (see Figure S6).

### CellSignals

One of `gro’`s hallmarks is the simulation of environmental signals. Externally inoculated signals like chemical inducers (such as arabinose, IPTG, or aTc) enable or disable the expression of proteins. The most frequently studied form of intercellular communication with signals is quorum sensing (QS).(*43–45*) QS is a mechanism in which cells emit small diffusible molecules into the environment which are then read by other cells. Through absorption of the molecules, cells can quantify their density in terms of the concentration of molecules in the environment. QS has been extensively used in Synthetic Biology for intercellular communication. Examples of QS-based systems include pattern formation,^46,47^ population control,^48^ bacterial ecology^49^ and computing logic gates^50^ among other applications.

Signals in the original version of `gro` are implemented through a set of grids (one for each signal) that store the signal concentration at each grid location. At each time step of the simulation, diffusion and degradation are applied to update the concentrations of the signals over the whole grid. One limitation we found with the implementation is that the grids have a static size. This implies that the programmer must have a good idea of what size the colony will reach in the simulation, which is not always the case. Another important shortcoming is that `gro` uses a finite element model^51^ for signal diffusion and degradation with fixed coefficients. Since the user cannot introduce any parameters related to the diffusion method, the variety of experiments that can be specified using signals is limited.

In order to improve the signal implementation, we reorganized the structure, extended and encapsulated its functionality into a new external library called CellSignals. This library preserves the signal diffusion and degradation methods used in the original implementation. It also provides the option of customizing the diffusion method by tuning the coefficients that govern how the signal expands through the grid. It also provides a dynamic-sized grid that automatically resizes whenever it is needed (see Algorithm S8).

Along with the extension of CellSignals, we included some signal-related *actions* (see Table 1). Absorption of molecules can now trigger the expression of binary proteins or modulate growth. These dynamics seek to simulate QS and cross-feeding mechanisms respectively. The QS mode of absorption checks for a given amount of signal concentration, and consumes this signal from the environment. If the amount of absorbed signal is over a given threshold set by the user, it triggers the expression of a binary protein. This protein then integrates into the rest of the protein specification handled by CellPro. In the crossfeeding mode, a given amount of signal is also consumed from the environment. However, the effect of absorbing the signal in this case affects the growth rate of the bacterium. If the interaction type is set to positive (+), then bacteria would need to absorb a given amount of signal to grow normally: The amount of absorbed signal directly modulates the bacterial growth rate. On the other hand, if the interaction type is set to negative (−), bacteria are still forced to absorb, but now the signal effect is detrimental, slowing down the growth of the cell.

### Bacterial conjugation

Cell-cell communication mechanisms are key in coordinating tasks in bacterial communities. The two main types of communication methods are bacterial signaling, such as QS, and horizontal gene transfer (HGT). HGT are a class of short-range communication methods that exchange large amounts of information in the form of DNA to nearby cells.^52^ One example of an HGT process is bacterial conjugation.^53^

IbMs are best suited to simulate communication mechanisms since the transfer of information highly depends on the spatial arrangement of the individuals. `gro` natively implements intercellular signaling as a communication method. However, no other intercell communication method is present. Conjugation proves to be a programmable local communication system for bacteria,^16^ making it a valuable synthetic tool. Also, the inclusion of conjugation provides a platform for studying and understanding HGT processes. We are unaware of any other IbM simulator implementing more than a single intercelluar communication mechanism to date.

The conjugation method is programmed as an *action* triggered by the involved proteins. Its specification is done trough ProSpec (Table 1). When the conjugation *action* is triggered, it transfers a plasmid to a random neighbor bacterium at a specified conjugation frequency, *γ*. The logic for the conjugation algorithm is summarized in Algorithm S7.

In order to complement and add realism to the behavior of conjugation, we added some plasmid-related *actions*: plasmid segregation and entry exclusion. A transferred plasmid may not be able to survive within the receiving cell. It happens mainly because another plasmid of the same family is already residing in the receiving cell. This mechanism is called entry exclusion.^54^ A plasmid may also be lost upon division. Plasmid segregation is specially critical when a plasmid has a low copy number.^55^ Even though conjugation is not well understood, we believe that the presented plasmid operations can reproduce most scenarios involved in bacterial conjugation.

## Systems biology studies and synthetic biology examples

To illustrate the capabilities and flexibility of the new libraries we present two systems biology case studies: plasmid propagation and bacterial ecological interactions. These case studies show that `gro` can be a useful tool for research fields such as microbial evolutionary biology, ecology or epidemiology. In addition, we show two brand-new synthetic circuits. All examples were simulated with the new version of `gro`. Experiment descriptions and videos can be found in the supporting information.

In the first case study we show how conjugation is implemented. We enquired the factors affecting conjugative plasmid propagation in a surface-associated bacterial population.^56^ Specifically, we studied the influence of space clumping for the infection to thrive. Results are shown in Figure 6.

**Figure 6:**
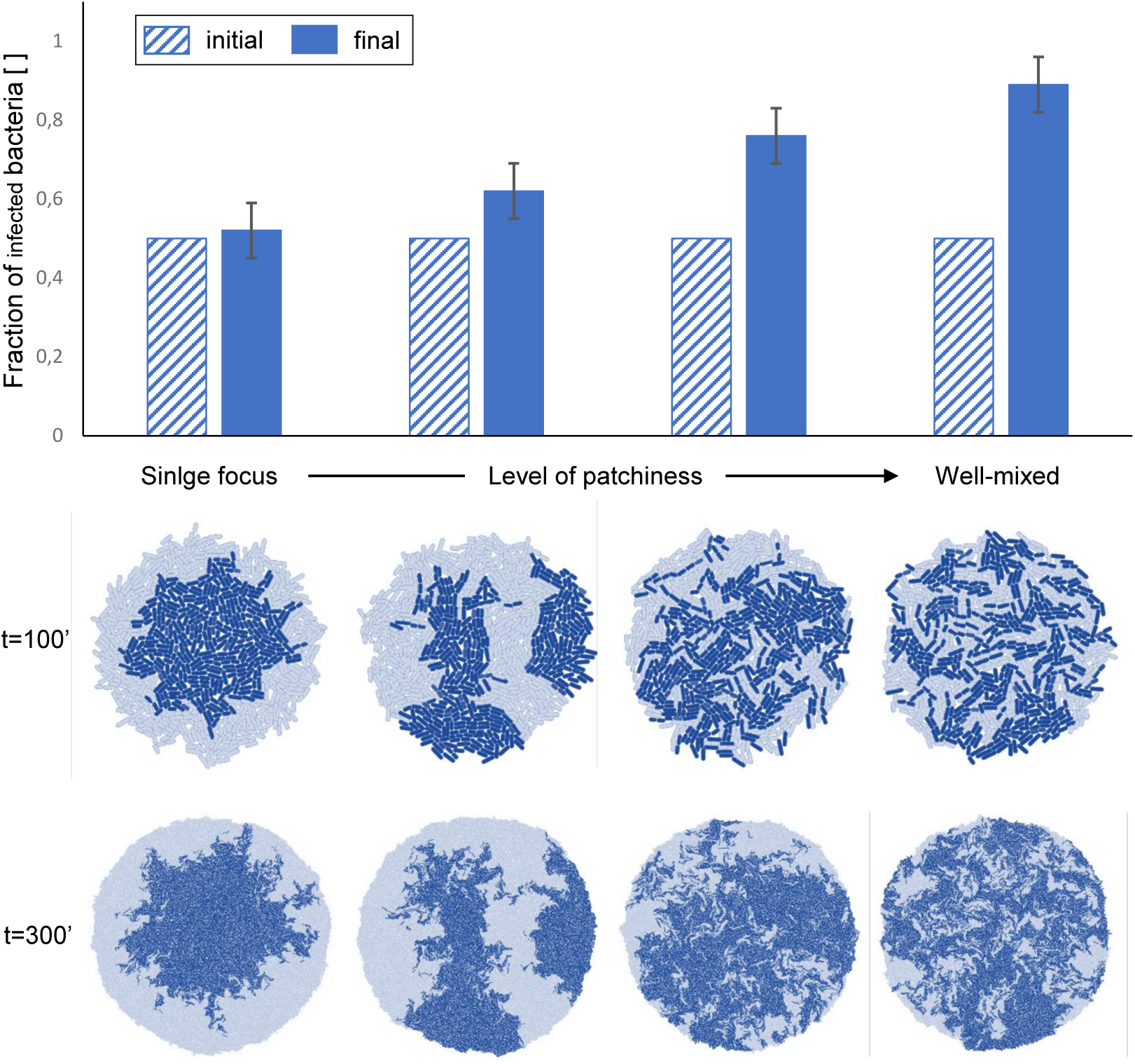
Spatial clumping affects plasmid propagation. Simulations were run for four different spatial arrangements ranging from a single focus (maximal patchiness) to a well-mixed case (minimal patchiness). All experiments shared the same conditions: half of the initial population was infected with a conjugative plasmid that transferred at a conjugation frequency of 0.3 while the other half was susceptible to the infection. Plasmid-bearing cells had a generation time of roughly 44 minutes while plasmid-free cells divided around every 40 minutes. Results are shown in the bar graph. Each bar corresponds to the spatial arrangement right below. Error bars display data corresponding to five runs. As it can be observed, the final fraction of infected bacteria drops from a 0.9 in the well-mixed case, to a 0.5 for the single focus scenario.

A critical factor affecting plasmid propagation is the ability of the infected cells to contact uninfected ones. In Figure 6 we present results from the plasmid propagation experiments and its relation with different spatial initial conditions. The simulation considers plasmid-free bacteria and plasmid-bearing bacteria that grow and interact trough physical contact. All experiments were run under the same conditions, apart from the initial spatial distribution. Half of the initial population are plasmid-bearing bacteria that carry an extra metabolic burden imposed onto them as they must conjugate. Plasmid conjugation frequency *γ* was fixed at 0.3 conjugative events during the cell life cycle, which was set to 44 minutes. After 300 minutes of experiment time, resulted colonies were analyzed. As it can be observed in Figure 6, the infected population fraction drops from a 0.9 in the well-mixed case, to a 0.5 for the single focus scenario. Results are consistent with the basic condition for plasmid propagation: large contact surface between plasmid-free and plasmid-bearing cells improves plasmid spread trough conjugation. Still, it is worth noticing that clumping has a dramatic effect which should not be overseen in precise computations of plasmid invasion.^57^

To further illustrate the new capabilities of the software, the second case study drives along the lines of pattern formation. Bacterial ecological interactions are tested in simulations in which two strains either compete or cooperate through cross-feeding signals. In this setup, diffusive molecules are emitted by bacteria themselves, and their effect on the opposite strain results in diverse spatial patterns which are shown in Figure 7. Three ecological interactions were programmed for a two-strain consortium. The neutralism case represents a colony of two types of bacteria that interact only through mechanical contact. In the cooperative consortium, bacteria emit a signal which is required by their counterpart to grow appropriately. The absorption of the molecule induces growth which in turn triggers the emission of the outgoing bacterial signal. The requirement of each other force them to grow as closer as possible. Thus, inter-mixing patterns arise in this particular case.^58^ On the other hand, when the absorption of a molecule is set to be detrimental for the other, bacteria repel each other as their emissions are toxic and prevent growth. In this scenario, the contact surface between the two strains is minimized, leading in some cases to the total annihilation of one of the strains. Studies of this kind can be further elaborated to include a variety of ecological behaviors between two or more strains^59,60^

**Figure 7:**
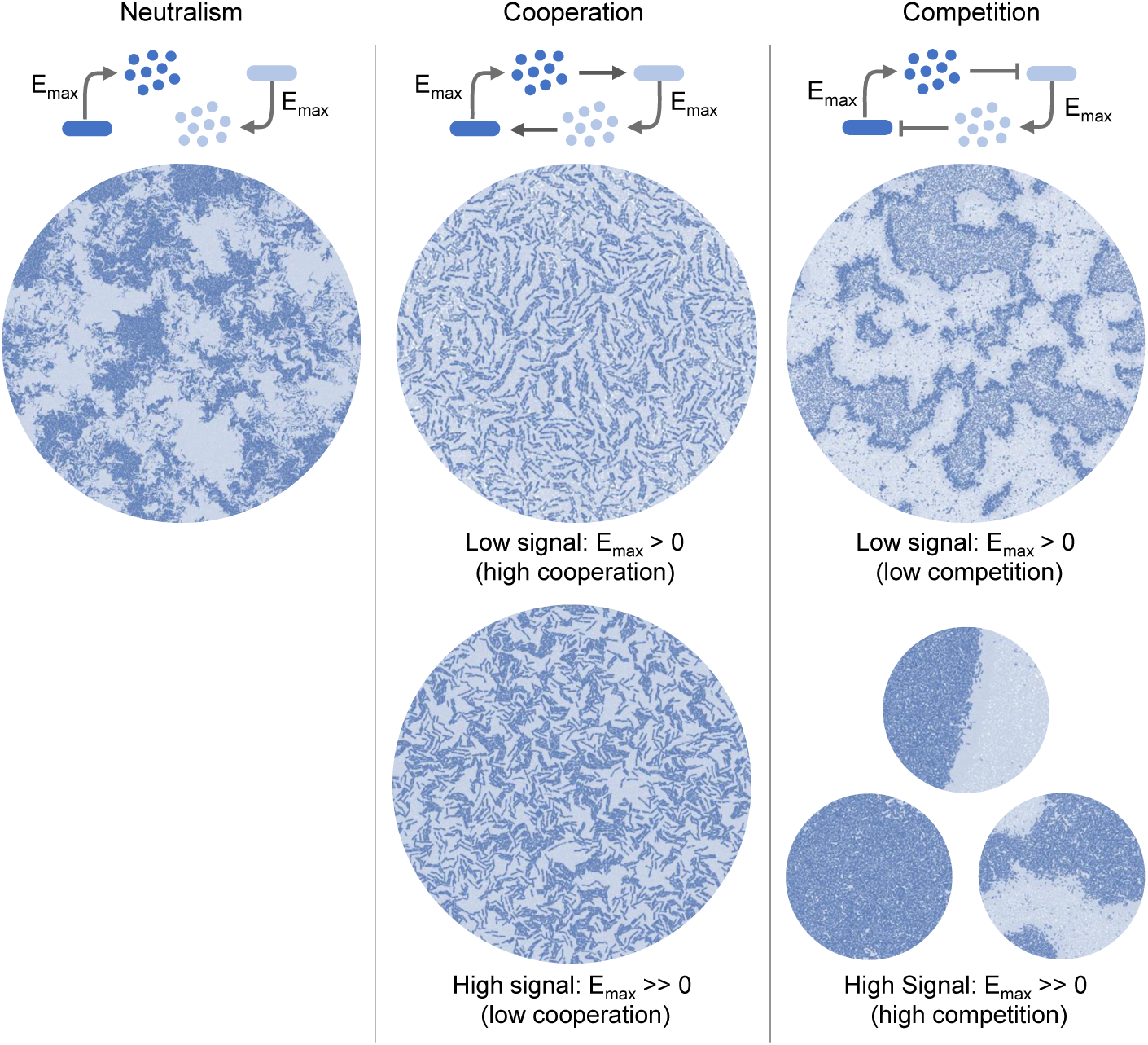
Ecological cross-feeding interactions and their resulting spatial patterns. Three ecological interactions were programmed for a two-strain consortium. The neutralism case represents a colony of two types of bacteria that interact only through mechanical contact. In this case the spatial pattern is created only by the random distribution of growing bacteria. When the interaction type is set to cooperation, bacteria emit a signal which is required by their counterpart to grow and emit its signal appropriately. The emission rate *E_max_* is set externally, leading to different interaction strengths. When the emission rate *E_max_* is low, molecule concentration in the environment is limited, and cooperation becomes forced. The outcome is a pattern of highly mixed strains that try to maximize access to the molecule. When the emission rate *E_max_* is high, the abundance of signal does not force bacteria to grow as close and the spatial patterns tend to the neutralism case. In the competition case, the logic is similar to a toxin emission that only affects the counterpart strain. Such interaction results in the avoidance of the other strain and bacteria arrange in clusters to minimize contact surface. However, when toxin concentration is in shortage, the toxicity in the medium is low and coexistence is possible. Nevertheless, bacteria at the interface will not grow nor emit. In the case of environmental signal abundance, the toxicity rises leading to a strong competition, which is highly sensitive to initial conditions. Images from different runs are shown proving that many scenarios are possible ranging from inhomogeneous coexistence to total annihilation of one of the strains.

Apart from the two exposed systems biology experiments, we also designed and simulated two original synthetic circuits based on cell-cell communication. In the first design, a three level bullseye pattern is generated based on a simple diffusible environmental signal (IPTG) as template (see Figure 8a). This first circuit is composed of three conjugative plasmids distributed equally within the initial population. Only two of the three plasmids are sensitive to the presence of IPTG. In these two plasmids, reporting is respectively induced and repressed under IPTG, discriminating the space into high and low IPTG levels. An intermediate diffusible signal is expressed by both plasmids in order to repress the reporting generated by the third plasmid. This dynamics, along with a low metabolic cost of the third plasmid result in the concentric alignment of the three synthetic strains shown in Figure 8a. In all cases, plasmid conjugation and reporting are regulated together, reinforcing the plasmid presence wherever it reports.

**Figure 8:**
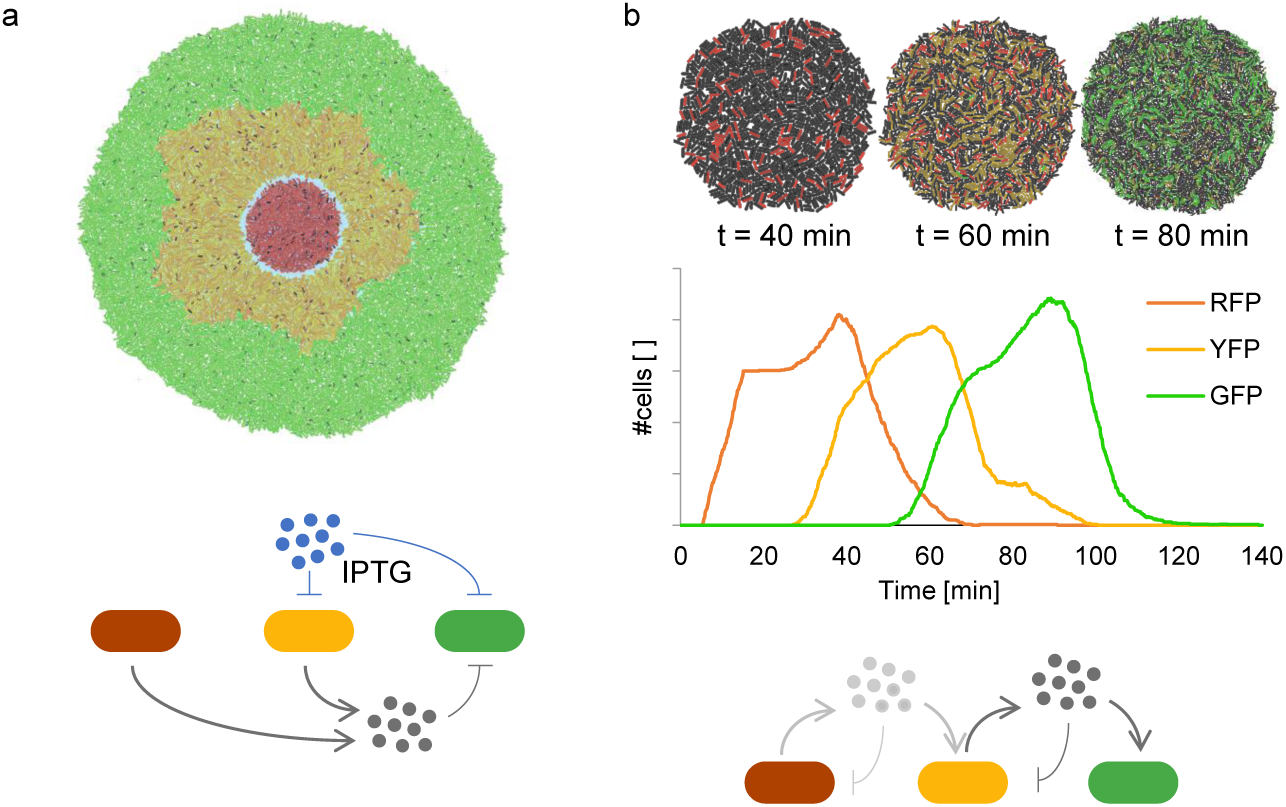
Spatial and temporal patterns simulated with `gro`. a) A three-level bullseye spatial pattern is depicted. The pattern described by the cells arises from processing a diffusible IPTG environmental signal as template (in blue). IPTG is found in higher concentration in the center of the colony and decreases radially. Cells in presence of a high concentration of IPTG express RFP and emit AHL. A middle crown is formed by cells expressing YFP that cannot survive in a zone where there is a high concentration of IPTG. These cells also emit AHL (gray). An outer crown of GFP expressing cells results from the expression of a toxin (triggered by the detection of AHL) that kills them and drives them towards the edge. In summary, IPTG eliminates YFP and GFP expressing cells from the center of the colony, and AHL forces GFP cells to grow on the outer part of the colony by triggering their death in the inner part of the bacterial population. The shown pattern is obtained after 300 simulated minutes. b) A signal based circuit for producing sequential protein expression pulses at a colony-level is shown. The circuit uses intercellular signals to guide the colony to dominantly express one of the three fluorescent proteins in a sequential order. Each signal has a double purpose: to trigger the destruction of its emitting plasmid and to activate the next pulse. Pulses reach their peak at 37 minutes (RFP), 61 minutes (YFP) and 89 minutes (GFP). Further details and both design specifications can be found in Figure S8.

The last synthetic circuit we present here is a distributed circuit that sequentially expresses three different proteins (reporting proteins in this case) in a specific order and for a given duration. Each step of the sequence translates to the expression of a single protein which is encoded in a plasmid. In this way only bacteria bearing the applicable plasmid will express the corresponding protein. A cell can only execute one step of the sequence, since no cell bears more than one of the three implicated plasmids. Each plasmid encodes for a different diffusible signal that triggers the start of the next step of the sequence in nearby cells. This same signal initiates the destruction of the signaling plasmid, stopping the current step of the sequence through a delayed negative feedback. We depict the design and results obtained for this circuit in Figure 8b. Although it is only a proof of concept, the idea behind this design is to mimic a well mixed bioreactor where every one of the agents is specialized in one stage of the path.^61^ Crosstalk and inefficiency are avoided by inducing the different steps of the path sequentially. In addition, every cell is involved in a single step of the sequence, allowing every step to be more complex and metabolically heavy for the host. The presented design is scalable, making it feasible to arrange larger sequential paths.

## Discussion

We presented a new version of `gro`, an open-source fast prototyping tool for the simulation of microbial multicellular colonies. The presented software is useful for studying and simulating complex intercellular processes. This `gro` version executes between one and two orders of magnitude faster than the original version and around an order of magnitude faster than other proficient IbM simulators like CellModeller. The modifications made to `gro` seek to improve the performance of the simulator and to extend its capabilities to make it attractive and accessible for a broad audience coming from multiple fields such as synthetic biology, systems biology, evolutionary microbiology or epidemiology.

As a proof of concept, we show comprehensive and easy to implement examples related to these diverse fields. Each example exhibits a high degree of precision within the scope of its theme, proving that `gro` is a versatile tool.

We believe that this new version of `gro` can overcome performance barriers, such as computing speed and colony size.^23^ We expect `gro` to evolve and become even more intuitive and scalable. Once it evolves, more realistic simulation of synthetic biology circuits will be achieved. At that point, analysis of multicellular systems biology problems and microbial ecosystems will bear better and more accurate results.

## Future work

The modular architecture of `gro` enables enables a relatively straightforward attachment of additional biological tools. We foresee the inclusion of more tools such as phages, CRISPR and RNA regulation, which are at present under development in our lab.

We are also currently looking at other platforms onto which the application of CellEngine could boost their performance. Other types of bacteria with different shapes, like yeast, are being considered as an extension to CellEngine. We plan to make CellNutrient a part of CellSignals, as nutrients may be treated as a specific case of signal. We are also currently researching on diffusion methods using binary valued signals to extend the capabilities of CellSignals.

Direct import/export of the models is another interesting direction: Use of SBOL 2.0^62^ enriched with quantitative modeling data in the form of annotations could serve as the basis of a standard description language for IbM simulations. Additionally, a graphical user interface (GUI) for assisting in experiment specification is being developed. We wish to minimize human error when specifying an experiment and to provide an intuitive and easy way for manipulating simulations. These are natural improvements, since ProSpec was designed keeping genetic circuit structure in mind.

## Material and Methods

CellEngine and CellNutrient were implemented in C as external libraries that integrate with `gro`. CellSignals and CellPro were implemented in C++ and are also external libraries that integrate with `gro`. Conjugation was implemented as a direct extension to the source code of `gro` in C++. The latest version of QT Creator framework (5.7) was used as the IDE to implement these features. Both the external libraries and the modified `gro` version were tested on Linux CentOS 6.6 and 7.0, and on MacOS X 10.10, 10.11 and 10.12. All examples were run on a Mac Mini Quad core i7, 2.1 GHz with 16 GB RAM and on Intel based Quad core machines 2.5 GHz with 32 GB RAM. The source code for the simulator and all of the simulations may be found at: http://www.lia.upm.es/software/gro/

## Acknowledgement

This work was supported by the European Union project PLASWIRES (612146/FP7-ICT-FET-Proactive) and by spanish MINECO projects TIN2012-36992 and TIN2016-81079-R. This work was also supported by Universidad Diego Portales and CONICYT Chile through the Becas Chile program.

## References

1. Macía, J., Posas, F., and Solé, R. V. (2012) Distributed computation: the new wave of synthetic biology devices. Trends Biotechnol. 30, 342–349.

2. Gillespie, D. T. (1977) Exact stochastic simulation of coupled chemical reactions. J. Phys. Chem. 81, 2340–2361.

3. Karlebach, G., and Shamir, R. (2008) Modelling and analysis of gene regulatory networks. Nat. Rev. Mol. Cell Biol. 9, 770–780.

4. Hellweger, F. L., Clegg, R. J., Clark, J. R., Plugge, C. M., and Kreft, J.-U. (2016) Advancing microbial sciences by individual-based modelling. Nat. Rev. Microbiol. 14, 461–471.

5. Gorochowski, T. E. (2016) Agent-based modelling in synthetic biology. Essays Biochem. 60, 325–336.

6. Song, H.-S., Cannon, W., Beliaev, A., and Konopka, A. (2014) Mathematical Modeling of Microbial Community Dynamics: A Methodological Review. Processes 2, 711–752.

7. Rudge, T. J., Steiner, P. J., Phillips, A., and Haseloff, J. (2012) Computational modeling of synthetic microbial biofilms. ACS Synth. Biol. 1, 345–352.

8. Lardon, L. A., Merkey, B. V., Martins, S., Dötsch, A., Picioreanu, C., Kreft, J.-U., and Smets, B. F. (2011) iDynoMiCS: next-generation individual-based modelling of biofilms. Environ. Microbiol. 13, 2416–2434.

9. Kang, S., Kahan, S., McDermott, J., Flann, N., and Shmulevich, I. (2014) Biocellion: accelerating computer simulation of multicellular biological system models. Bioinformatics 30, 3101–3108.

10. Georgiev-Lab, CeCe. https://github.com/GeorgievLab/CeCe/, Accessed December 30, 2016.

11. Jang, S. S., Oishi, K. T., Egbert, R. G., and Klavins, E. (2012) Specification and simulation of synthetic multicelled behaviors. ACS Synth. Biol. 1, 365–374.

12. Gorochowski, T. E., Matyjaszkiewicz, A., Todd, T., Oak, N., Kowalska, K., Reid, S., Tsaneva-Atanasova, K. T., Savery, N. J., Grierson, C. S., and di Bernardo, M. (2012) BSim: an agent-based tool for modeling bacterial populations in systems and synthetic biology. PLoS One 7, e42790.

13. García, A. P., and Rodríguez-Patón, A. BactoSim-An Individual-Based Simulation Environment for Bacterial Conjugation. International Conference on Practical Applications of Agents and Multi-Agent Systems. 2015; pp 275–279.

14. Krone, S. M., Lu, R., Fox, R., Suzuki, H., and Top, E. M. (2007) Modelling the spatial dynamics of plasmid transfer and persistence. Microbiology 153, 2803–2816.

15. Harcombe, W. R., Riehl, W. J., Dukovski, I., Granger, B. R., Betts, A., Lang, A. H., Bonilla, G., Kar, A., Leiby, N., and Mehta, P. (2014) Metabolic resource allocation in individual microbes determines ecosystem interactions and spatial dynamics. Cell Rep. 7, 1104–1115.

16. Goñi-Moreno, A., Amos, M., and de la Cruz, F. (2013) Multicellular computing using conjugation for wiring. PLoS One 8, e65986.

17. Collier, N. (2003) Repast: An extensible framework for agent simulation. The University of Chicagos Social Science Research 36, 2003.

18. Kiran, M., Richmond, P., Holcombe, M., Chin, L. S., Worth, D., and Greenough, C. FLAME: simulating large populations of agents on parallel hardware architectures. Proceedings of the 9th International Conference on Autonomous Agents and Multiagent Systems: Volume 1. 2010; pp 1633–1636.

19. Tisue, S., and Wilensky, U. Netlogo: A simple environment for modeling complexity. International Conference on Complex Systems. 2004; pp 16–21.

20. Luke, S., Cioffi-Revilla, C., Panait, L., Sullivan, K., and Balan, G. (2005) Mason: A multiagent simulation environment. Simulation 81, 517–527.

21. Klavins, E. GRO Github. https://github.com/klavinslab/gro, Accessed December 30, 2016.

22. Solé, R., Amor, D. R., Duran-Nebreda, S., Conde-Pueyo, N., Carbonell-Ballestero, M., and Montañez, R. (2016) Synthetic collective intelligence. Biosystems 148, 47–61.

23. Pascalie, J., Potier, M., Kowaliw, T., Giavitto, J.-L., Michel, O., Spicher, A., and Doursat, R. (2016) Developmental Design of Synthetic Bacterial Architectures by Morphogenetic Engineering. ACS Synth. Biol. 5, 842–861.

24. Klavins, E. GRO simulation after five days. https://www.youtube.com/watch?v=P0ykJZhcOwI, Accessed December 30, 2016.

25. Lembcke, S., and Software, H. M. Chipmunk 2D Physics. https://chipmunk-physics.net/, Accessed December 30, 2016.

26. Klavins, E. A language for modeling and programming cooperative control systems. Proceedings of the IEEE International Conference on Robotics and Automation. 2004; pp 3403–3410.

27. Mirtich, B., and Canny, J. Impulse-based simulation of rigid bodies. Proceedings of the Symposium on Interactive 3D graphics. New York, New York, USA, 1995; pp 181–189.

28. Ericson, C. Real-time collision detection; CRC Press, 2004.

29. Millington, I. Game physics engine development; Morgan Kaufmann Publishers Amsterdam, 2007.

30. Appel, A. W. (1985) An efficient program for many-body simulation. SIAM J. Sci. Comput. 6, 85–103.

31. Chen, P.-N., Lin, H.-Y., and Moser, S. M. Ultra-small block-codes for binary discrete memoryless channels. Proceedings of the IEEE Information Theory Workshop. 2011; pp 175–179.

32. Stoelinga, M. (2002) An introduction to probabilistic automata. Bull. Eur. Assoc. Theor. Comput. Sci. EATCS 78, 2.

33. Kwiatkowska, M., Norman, G., Segala, R., and Sproston, J. (2002) Automatic verification of real-time systems with discrete probability distributions. Theor. Comput. Sci. 282, 101–150.

34. Lloyd, D. P. Microscopic studies of surface growing bacterial populations. Ph.D. thesis, 2015.

35. Volfson, D., Cookson, S., Hasty, J., and Tsimring, L. S. (2008) Biomechanical ordering of dense cell populations. Proc. Natl. Acad. Sci. U. S. A. 105, 15346–15351.

36. Sáez, S. Diseño e implementación de un módulo de crecimiento bacteriano dependiente de nutrientes en el simulador GRO. B.S. thesis, 2015.

37. Monod, J. (1949) The growth of bacterial cultures. Annu. Rev. Microbiol. 3, 371–394.

38. Pirt, S. J. (1967) A kinetic study of the mode of growth of surface colonies of bacteria and fungi. J. Gen. Microbiol. 47, 181–197.

39. Drazek, L., Tournoud, M., Derepas, F., Guicherd, M., Mahé, P., Pinston, F., Veyrieras, J.-B., and Chatellier, S. (2014) Three-dimensional characterization of bacterial microcolonies on solid agar-based culture media. J. Microbiol. Methods 109, 149–156.

40. Lewis, M. W., and Wimpenny, J. W. (1981) The influence of nutrition and temperature on the growth of colonies of Escherichia coli K12. Can. J. Microbiol. 27, 679–684.

41. Farrell, F. D. C., Hallatschek, O., Marenduzzo, D., and Waclaw, B. (2013) Mechanically driven growth of quasi-two-dimensional microbial colonies. Phys. Rev. Lett. 111, 1–8.

42. Tokita, R., Katoh, T., Maeda, Y., Wakita, J., Sano, M., Matsuyama, T., and Matsushita, M. (2009) Pattern Formation of Bacterial Colonies by Escherichia coli. J. Phys. Soc. Jpn. 78, 074005.

43. Fuqua, C., Parsek, M. R., and Greenberg, E. P. (2001) Regulation of gene expression by cell-to-cell communication: acyl-homoserine lactone quorum sensing. Annu. Rev. Genet. 35, 439–468.

44. Miller, M. B., and Bassler, B. L. (2001) Quorum sensing in bacteria. Annu. Rev. Microbiol. 55, 165–199.

45. Waters, C. M., and Bassler, B. L. (2005) Quorum sensing: cell-to-cell communication in bacteria. Annu. Rev. Cell Dev. Biol. 21, 319–346.

46. Tabor, J. J., Salis, H. M., Simpson, Z. B., Chevalier, A. a., Levskaya, A., Marcotte, E. M., Voigt, C. a., and Ellington, A. D. (2009) A synthetic genetic edge detection program. Cell 137, 1272–1281.

47. Basu, S., Gerchman, Y., Collins, C. H., Arnold, F. H., and Weiss, R. (2005) A synthetic multicellular system for programmed pattern formation. Nature 434, 1130–1134.

48. You, L., Cox, R. S., Weiss, R., and Arnold, F. H. (2004) Programmed population control by cell–cell communication and regulated killing. Nature 428, 868–871.

49. Balagaddé, F. K., Song, H., Ozaki, J., Collins, C. H., Barnet, M., Arnold, F. H., Quake, S. R., and You, L. (2008) A synthetic Escherichia coli predator–prey ecosystem. Mol. Syst. Biol. 4, 187.

50. Tamsir, A., Tabor, J. J., and Voigt, C. A. (2011) Robust multicellular computing using genetically encoded NOR gates and chemical ‘wires’. Nature 469, 212–215.

51. Douglas, J., Jr, and Russell, T. F. (1982) Numerical methods for convection-dominated diffusion problems based on combining the method of characteristics with finite element or finite difference procedures. SIAM J. Numer. Anal. 19, 871–885.

52. Ortiz, M. E., and Endy, D. (2012) Engineered cell-cell communication via DNA messaging. J. Biol. Eng. 6, 16.

53. Smillie, C., Garcillán-Barcia, M. P., Francia, M. V., Rocha, E. P. C., and de la Cruz, F. (2010) Mobility of plasmids. Microbiol. Mol. Biol. Rev. 74, 434–452.

54. Garcillán-Barcia, M. P., and de la Cruz, F. (2008) Why is entry exclusion an essential feature of conjugative plasmids? Plasmid 60, 1–18.

55. Gordon, S., Rech, J., Lane, D., and Wright, A. (2004) Kinetics of plasmid segregation in Escherichia coli. Mol. Microbiol. 51, 461–469.

56. Zhong, X., Droesch, J., Fox, R., Top, E. M., and Krone, S. M. (2012) On the meaning and estimation of plasmid transfer rates for surface-associated and well-mixed bacterial populations. J. Theor. Biol. 294, 144–152.

57. Magal, P., and Ruan, S. (2014) Susceptible-infectious-recovered models revisited: from the individual level to the population level. Math. Biosci. 250, 26–40.

58. Momeni, B., Brileya, K. A., Fields, M. W., and Shou, W. (2013) Strong inter-population cooperation leads to partner intermixing in microbial communities. eLife 2013, 1–23.

59. Blanchard, A. E., and Lu, T. (2015) Bacterial social interactions drive the emergence of differential spatial colony structures. BMC Syst. Biol. 9, 59.

60. Reichenbach, T., Mobilia, M., and Frey, E. (2007) Mobility promotes and jeopardizes biodiversity in rock-paper-scissors games. Nature 448, 1046–1049.

61. Widder, S. et al. (2016) Challenges in microbial ecology: building predictive understanding of community function and dynamics. ISME J. 10, 2557–2568.

62. Roehner, N. et al. (2016) Sharing Structure and Function in Biological Design with SBOL 2.0. ACS Synth. Biol. 5, 498–506.

63. Bonnet, J., Subsoontorn, P., and Endy, D. (2012) Rewritable digital data storage in live cells via engineered control of recombination directionality. Proc. Natl. Acad. Sci. U. S. A. 109, 8884–8889.

64. Scott, S. R., and Hasty, J. (2016) Quorum Sensing Communication Modules for Microbial Consortia. ACS Synth. Biol. 5, 969–977.

65. Lakshmi, O. S., and Rao, N. (2009) Evolving Lac repressor for enhanced inducibility. Protein Eng., Des. Sel. 22, 53–58.

